# High-throughput characterisation of bull semen motility using differential dynamic microscopy

**DOI:** 10.1101/388777

**Authors:** Alys Jepson, Jochen Arlt, Jonathan Statham, Mark Spilman, Katie Burton, Tiffany Wood, Wilson C. K. Poon, Vincent A. Martinez

**Affiliations:** SUPA, School of Physics & Astronomy, The University of Edinburgh, Peter Guthrie Tait Road, Edinburgh EH9 3FD, UK; School of Biosciences, University of Exeter, Exeter, UK; RAFT Solutions Ltd., Mill Farm, Studley Road, Ripon, HG4 2QR, UK

## Abstract

We report a high-throughput technique for characterising the motility of spermatozoa using differential dynamic microscopy. A large field of view movie (~ 10mm^2^) records thousands of cells (e.g. ≈ 5000 cells even at a low cell density of 20 × 10^6^ cells/ml) at once and yields averaged measurements of the mean (*υ*) and standard deviation (*σ*) of the swimming speed, a head oscillation amplitude (*A*_0_) and frequency (*f*_0_), and the fraction of motile spermatozoa (*α*). Interestingly, the measurement of *α* relies on the swimming spermatozoa enhancing the motion of the non-swimming population. We demonstrate the ease and rapidity of our method by performing on-farm characterisation of bull spermatozoa motility, and validate the technique by comparing laboratory measurements with tracking. Our results confirm the long-standing theoretical prediction that 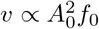 for swimming spermatozoa.

## Introduction

Sexual reproduction in all metazoans relies on the fertilisation of an ovum (egg) by a motile spermatozoon, which has to migrate through a variety of external or internal liquid environments to reach its destination. Motility is therefore of the essence of spermatozoon function, and the description of motile spermatozoa went back to the earliest days of scientific microscopy [1].

Spermatozoon phenotype is hugely variable across different phyla, both in terms of morphology and swimming characteristics, possibly as a result of co-evolution with the female reproductive tract [2]. Significant variability remains within the single subphylum Vertebrata. Thus, major adaptations were needed in the spermatozoon when marine vertebrates relying on external fertilisation evolved into terrestrial dwellers reproducing by internal fertilisation [3]. In both cases, the composition and properties of the seminal fluid in which spermatozoa are released help determine reproductive success [4].

Characterisation of spermatozoon motility in the male ejaculate is practiced as a crucial part of fertility assessment in humans as well as farm animals. Commercial bull semen evaluations are part of both routine pre-breeding examination (on-farm) with natural service and of monitoring the detrimental effect of storing, transporting and defrosting straws (in-lab) for artificial insemination (AI). The actual trajectory of the head of a bull spermatozoa is complex (Fig. 1 shows an example). For practical purposes, on farm, high quality motility is usually associated with a *high enough fraction* of spermatozoa showing *high enough progressive motility*, where the latter is associated with swimming along a straight trajectory. The two italicised quality factors are typically assessed visually (in a microscope) by an expert, with large margins of uncertainty. Thus, e.g., on-farm visual assessments of bull semen are found to have a variation of 20-40% [5].

**Fig 1.**
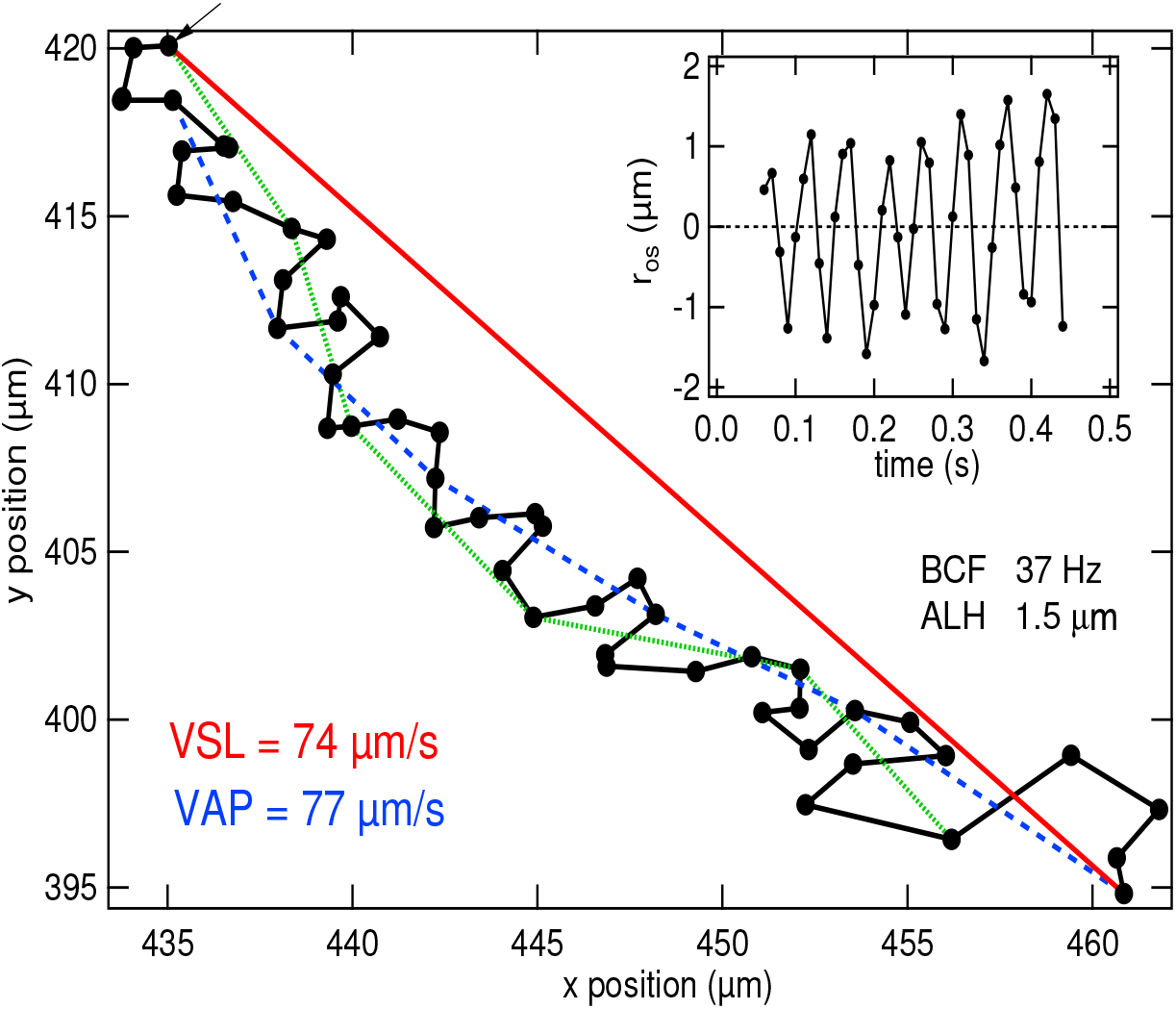
**Example of tracking a spermatozoon head motion,** using custom-made software (see Materials and Method for details) based on a movie recorded at 10 × magnification at 100fps for 0.5 s. VSL (red),VAP (blue and green), VCL (black), ALH and BCF calculated for this track are stated (see text for the definitions). Coloured lines indicate the distance used in calculating the corresponding speed (see section on Tracking analysis for explanation of VAP). Inset: The oscillatory component of displacement *r_os_* plotted against time.

**Fig 2.**
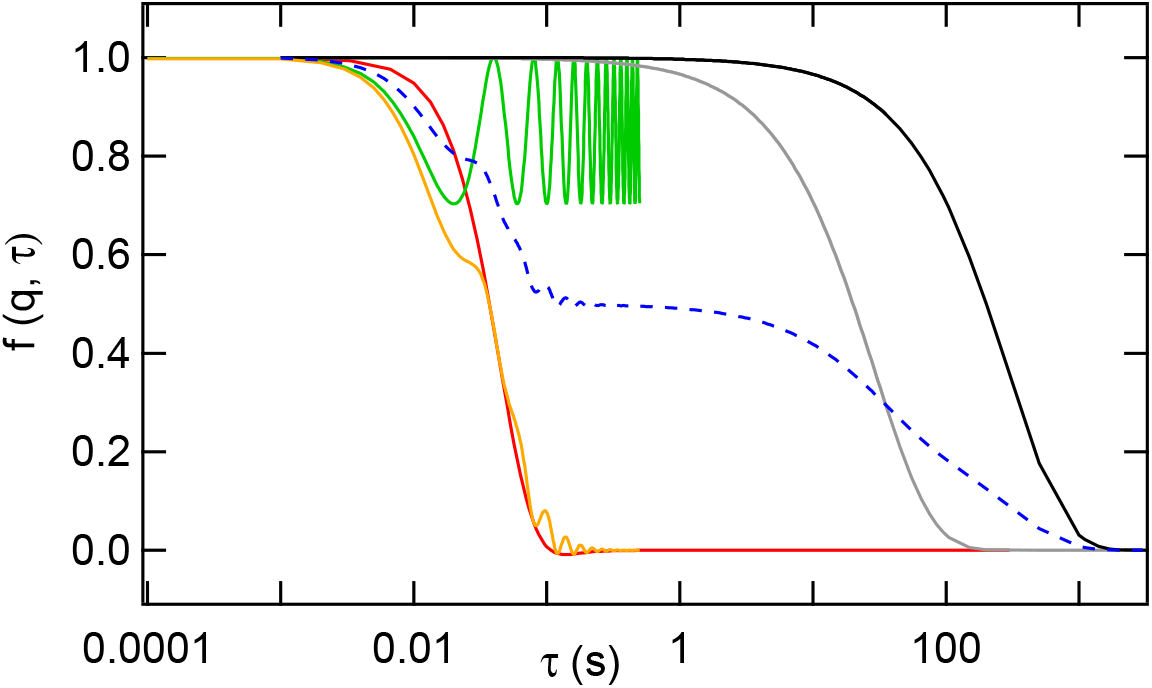
**Examples of theoretical** *f*(*q, τ*) at *q* = 0.34*μ*m^-1^ for typical parameters. *f*_m_(*q, τ*) from Eq. 7 for oscillatory motion (green): *v* = *σ* = 0 μms^-1^, *f*_0_ = 25Hz, *A*_0_ = 3μm; ballistic motion (red): *v* = 150μms^-1^, *σ* = 70μms^-1^, *f*_0_ = 0Hz, *A*_0_ = 0μm; oscillatory and ballistic motion (orange): *v* = 150μms^-1^, *σ* = 70μms^-1^, *f*_0_ = 25Hz, *A*_0_ = 3μm; diffusive motion (grey): *f*_d_(*q, τ*) for *D*_d_ = 0.3 μm^2^ s^-1^, and diffusive motion (black): *f*_nm_(*q, τ*) for *D*_nm_ = 0.03μm^2^ s^-1^. Full *f*(*q, τ*) (blue dotted): from all four (additive) contributions if swimming spermatozoa contribute 50% of the signal, debris 25% and non-motile spermatozoa 25%.

Computer-aided semen analysis (CASA) [6] can be used to give a more precise measure of the motile fraction, *α*, and to quantify motility. Spermatozoa typically swim by beating a single, flexible flagellum, causing the head to oscillate; near surfaces, they swim along curvilinear trajectories [7]. By tracking individual sperm cells and averaging over the population or more sophisticated sub-population analysis [8], CASA provides a number of kinematic parameters [9], including the actual-path velocity (VAP), the straight-line velocity (VSL), the head velocity calculated between successive frames (VCL), the amplitude of lateral head oscillations (ALH), and the beat cross frequency (BCF), Fig. 1.

CASA is laboratory (rather than clinic or farm) based, relatively costly, and involves dilution of semen to enable individual cells to be unambiguously tracked. Detailed quantification using CASA is only performed in a minority of cases and not commonly used for assessment of bulls examined pre-breeding or for investigation of poor performance in frozen AI semen, or natural bull service. There is therefore a paucity of data to enable unambiguous correlation between various motility measures and fertility. Moreover, the effect of commonly-used freezing and thawing processes used in AI and IVF is poorly quantified to date.

Here, we demonstrate a method based on differential dynamic microscopy (DDM) [10–12] for spermatozoa motility characterisation usable at the point of semen collection. The method is fast enough to yield the necessary quantity of data to inform future studies of the correlation of motility parameters with fertility and to quantify time-dependent effects of handling protocols (freeze/thaw, etc.).

We set up and validate our method in the context of bovine fertility, where spermatozoa motility is recognised to be a key component in semen evaluation [13]. We demonstrate that DDM is a high-throughput on-farm technique to measure population averaged values of *α*, VAP, ALH and BCF immediately after semen collection, and validate our method by comparing DDM with tracking measurements in the laboratory.

Our method should be important beyond the farm and fertility clinic. Quantifying swimming behaviour is a key component in the study of other aspects of spermatozoon biology such as the response of mitochondrial membrane potential to myoinositol [14], the role of Ca^2+^ in regulating flagella activity, hyper-activation [15,16] and chemotaxis [17—19]. Our technique should also impact fundamental active matter physics, where various aspects of spermatozoon swimming attract attention, e.g. collective motion [20], swimming mechanics [21], cooperation and competition between cells [22], and movement against flow [23] and along surfaces [24–26]. As an illustration, we use our extracted motility parameters from bull semen to investigate the relationship between the kinetic parameters *υ*, *A*_0_ and *f*_0_, and validate a long-standing theoretical prediction, that 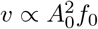.

## Materials and methods

### Theory

The principles of DDM have been described elsewhere [10–12,27]. Here we give a brief outline and apply the principles to spermatozoa. DDM uses low-resolution movies to obtain the differential image correlation function *g*(*q, τ*) (DICF), i.e. the power spectrum of the difference between pairs of images separated by delay time *τ*;

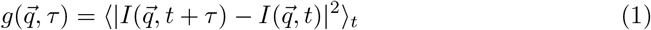

where 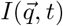 is the fourier transform of 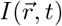, the intensity at pixel position 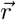 at time *t*, and the spatial frequency *q* = 2*π*/*l* defines the length scale *l* of interest. For isotropically moving cells, azimuthal averaging gives 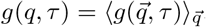. The DICF is related to the intermediate scattering function (ISF) *f*(*q*, *τ*) via [11]

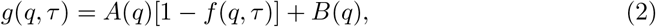

where *B*(*q*) accounts for instrumental noise. For non-interacting particles (here, swimming cells), *A*(*q*) ∝ *ϕa*(*q*) is the signal amplitude of particle population with *ϕ* the cell density and *a*(*q*) the signal amplitude of a single particle. The ISF *f*(*q, τ*) is related to the particle displacement Δ*r* by

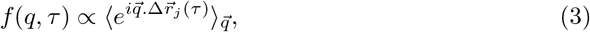

where ‘*j*’ denotes the *j*-th particle and brackets average over 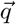 and all particles.

Semen samples contain two populations of spermatozoa (s), motile (m) and non-motile (nm), which have the same shape (head and flagellum) and hence *a*_m_(*q*) = *a*_nm_(*q*). Samples also contain debris (d) – particulates and/or cytoplasmic droplets – usually smaller than intact cells. We thus define the signal amplitude of the population *A_i_*(*q*) ∝ *ϕ_i_a_i_*(*q*) with ‘*i*’ = ‘s’ or ‘d’ for sperm cell or debris respectively. The *g*(*q, τ*) for such a semen sample is related to the ISFs for spermatozoa *f_s_*(*q, τ*) and for debris *f_d_*(*q, τ*) via

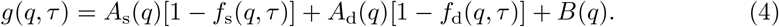

For a suspension containing motile 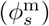 and non-motile 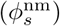 spermatozoa in proportions 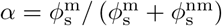 and 1 – *α* respectively,

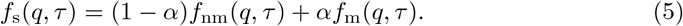

The ISF describes the decorrelation of particle positions with time and can be fitted with a theoretical model representing cell motion. Figure 1 suggests that we can model the movement of the sperm cell head using a linear progression with a superimposed sinusoidal motion:

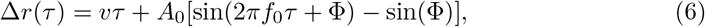

where *A*_0_ and *f*_0_ are the amplitude and frequency of head oscillation, Φ is a random phase and *v* is the swimming speed of linear progression. For non-interacting non-synchronized swimmers in 3D this returns [12],

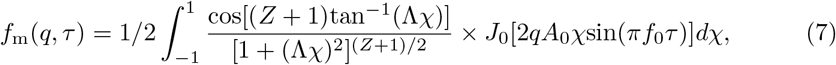

where Λ = *qvτ*/(*Z* + 1), *χ* = *cosψ* with *ψ* as the angle between 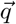 and 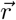 and *J*_0_ is the zeroth order Bessel function, assuming a Schultz distribution with a mean of *v* and a width of 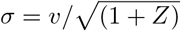 for the swimming speed distribution *P*(*v*). The same kind of function was used to extract swimming parameters from swimming algae [12].

We assume that the movement of non-motile spermatozoa and debris in a semen sample are diffusive, with diffusion coefficients *D*_nm_ and *D*_d_ respectively, so that

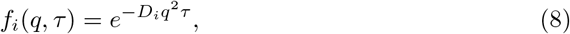

where the index *i* denotes either ‘nm’ or ‘d’.

In summary, we expect four contributions to *f*(*q, τ*) and hence to *g*(*q, τ*) for a semen sample: from head oscillation and ballistic motion of the swimmers and diffusion of non-motile spermatozoa and debris. Each contribution possesses a characteristic time scale, *t*_osc_, *t*_b_, *t*_*D*nm_ and *t*_*D*d_ respectively, which scales distinctly with *q* according to its motion: *t*_b_ ~ (*qv*)^-1^ (from the left term of the integrand in Eq. 7), *t*_osc_ = 1/*f*_0_ ~ *q*^0^ (from the *J*_0_ term in Eq. 7), and *t_D_* ~ (*q*^2^*D*)^-1^ (from Eq. 8). For motile sperm cells, the oscillatory and ballistic components are expected to crossover at *t*_osc_ = *t*_b_, where *q*_c_ ~ *f*_0_/*v*. The theoretical *f*(*q, τ*) calculated using a realistic set of bull spermatozoa parameters is plotted in Fig 2, together with the four separate contributions identified above.

Note that ISFs of bull spermatozoa [28–30] and other animal spermatozoa [31] were measured with dynamic light scattering in the 1970s. However, at the scattering vectors used (*q* ≥ 3.5μm^-1^), corresponding to length-scales *l* ≲ 1.8 μm, the signal was dominated by the oscillatory motion of the head, so that it proved impossible to obtain the swimming speed from fitting this data [30]. DDM overcomes this difficulty by accessing a wider range of length scales, over which swimming and head oscillation are well decoupled.

### Sample preparation, measurement and analysis

Some measurements were performed on fresh semen collected during a field study of bulls in South East Scotland undergoing routine breeding soundness examinations, approved by the Royal (Dick) School of Veterinary Studies (Veterinary Ethical Review Committee VERC Ref:29-14). Addition of phosphate-buffered saline (PBS) produced diluted samples. In other cases, we employed frozen semen used for artificial insemination from pooled Belgian Blue bulls (BB), a Holstein bull (HO) and a Charolais bull (CH) provided by RAFT Solutions Ltd. This frozen semen was contained in 0.25cc straws at a concentration of ≈ 80 ± 10 × 10^6^ cell/ml and stored in liquid nitrogen. Thawing of straws was performed in a 37 ^¤^C water bath for 30 s, after which the contents were immediately expelled into an Eppendorf tube using a metal rod. Samples at densities typically used for CASA (≈ (20 ± 10) × 10^6^ cell/ml) were obtained by diluting by 1:4 in Easybuffer B (IMV Technologies).

Samples were loaded into either 20 μm deep disposable counting chambers (Leja) or 50mm × 1 mm × 0.05 mm glass capillaries (VitroTubes), pre-warmed to 37 ¤C. The ends of chambers were sealed with vaseline to prevent drift. Samples were imaged in the centre of either kind of chamber.

Cell densities were estimated by manual counting from 10× micrographs, Fig. 3. The proportion of swimming spermatozoa was determined visually from movies, using ImageJ [32] to partition each frame into sectors and replaying the movie at a reduced frame rate (see S1 Fig).

**Fig 3.**
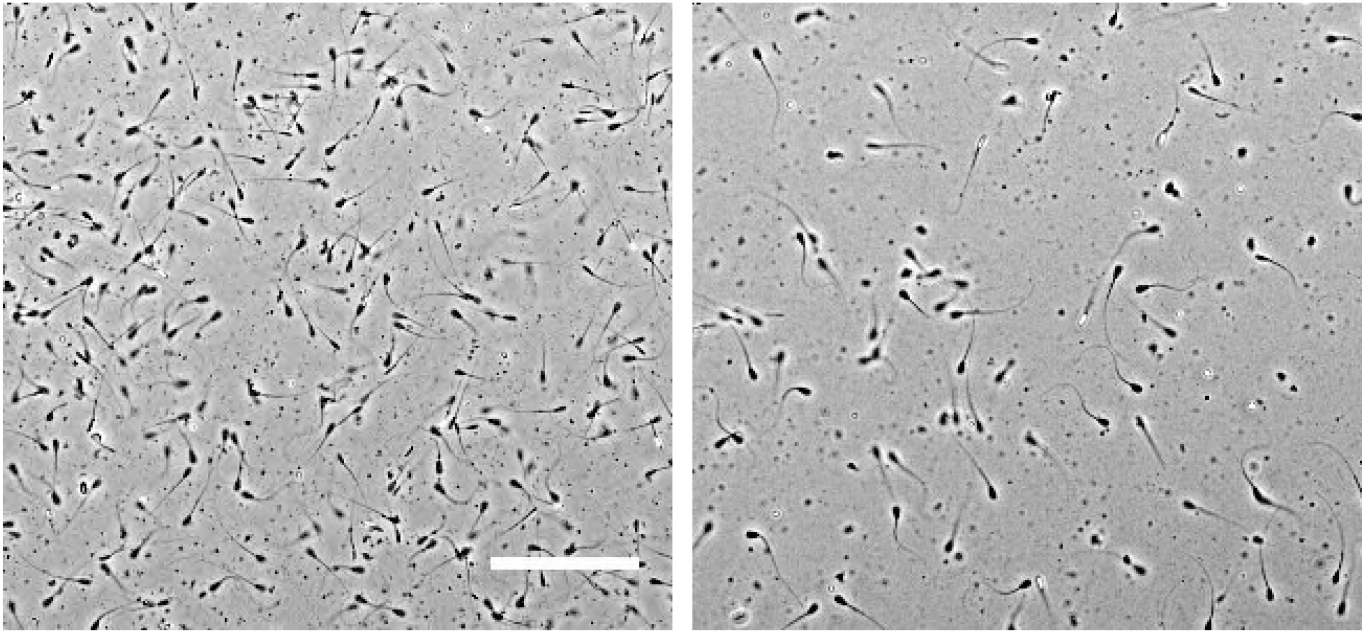
**Phase contrast micrographs** (10×) of (left) an undiluted thawed sample with ~ 80 × 10^6^ cell/ml and (right) a diluted thawed sample with ~ 20 × 10^6^ cell/ml. Scale bar=100 μm.

### DDM Analysis and Processing

Fresh samples were imaged on-farm using a home-made inverted microscope deploying a 2.5× Olympus objective and a uEye UI-1225LE-M-GL camera (IDS GmbH) giving an image with 2.65μm/pix. A Linkam MC60 heated stage was used to maintain the sample temperature at 37°C. Movies were recorded at 100 frames per second.

Samples thawed from frozen straws were imaged in the laboratory on a Nikon Eclipse Ti inverted microscope and recorded using a Mikrotron MC 1362 camera with a CMOS detector (pixel size 14 × 14μm^2^). The microscope was placed in an insulated box maintained at 37°C, where sample chambers were pre-warmed. To perform DDM, we recorded 2× (0.142 pix/μm) bright field movies (Nikon Plan Fluor, NA=0.06), or on occasions 10× phase contrast (NA=0.3) movies. Movies were recorded with frame sizes of 300 – 500 pixels at 100 or 300 frames per second, the movie length varying from ~ 5 – 100 s. In what follows we give the movie parameters in the format (magnification, framerate, frame size in pixels, movie length in seconds).

The DICFs, Eq. 1, were obtained from the movies by calculating the power spectrum of the difference between two images for a given delay time *τ* using custom-Labview software. These were averaged over a range of different initial times and scattering vectors 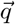. The calculation was repeated for a range of different τ to give *g*(*q*, *τ*). Further details of the calculation have been given elsewhere [12]. DDM processing and fitting analysis of a typical movie of 4000 images with 480 x 480 pixels takes just under 2 min.

### Tracking analysis

To perform particle tracking, 0.5s phase contrast movies at 10× magnification (Nikon Plan Fluor with NA=0.3) were obtained at 100 frames per second at a frame size of 500 – 1024 pixels. Standard tracking software [33] was used to obtain 2D trajectories, *r*(*t*), of spermatozoa head motion. All tracks were analysed to return the swimming parameters VSL= (*r*(0.5) – *r*(0))/0.5 and VAP = 〈(*r*(*t*) – *r*(*t* – 0.1))/0.1〉_*t*_, Fig. 1, where 〈…〉_*t*_ denotes averaging over time interval *t*. The time step of 0.1s was chosen as the shortest interval to return VAP=VSL for a straight track, while shorter time steps gave a speed that tended towards VCL. It was checked that the value of VAP obtained also corresponded closely to the speed along a path of 〈*r*〉_*dt*_ where *dt* = 2/*f*_0_.

To calculate *f*_0_ (corresponding to BCF/2) we analysed the oscillating component of displacement *r*_os_ = *r* – 〈*r*〉_*dt*_ in Fourier space [12]. The average head oscillation amplitude was calculated using 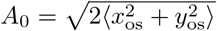, where *x_os_* = *x* – 〈*x*〉_*dt*_.

## Results & Discussion

We first describe results obtained on farm to demonstrate the portability and speed of our technique. Then we validate the technique in the laboratory by tracking.

### On-farm DDM

Figure 4(a) shows typical *g*(*q, τ*)’s measured from a movie of 25× diluted fresh semen in PBS with ~ 20 × 10^6^ cell/ml, from which we calculated *f*(*q, τ*). We first analysed this data by direct visual inspection and draw a number of order of magnitude conclusions. Then we fitted this data to obtain quantitative estimates of various motility parameters with associated error bars.

### Order of magnitude estimates

It is clear that *g*(*q, τ*) grows (and therefore *f*(*q, τ*) decays) in three steps, with three well-separated time scales, *t*_1_, *t*_2_ and *t*_3_, indicated by the three numbered arrows in Fig. 4(a). The fastest process is completed by *t*_1_ ≈ 0.06 s, and is *q*-independent. We identify this with head oscillations at 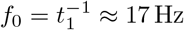. The intermediate time scale *t*_2_ of process 2 is *q*-dependent. Plotting the *f*(*q, τ*) calculated from *g*(*q, τ*) against *qτ* brings about collapse of the data for this process, Fig. 4(b), which is therefore ballistic. We read off a characteristic time at *qτ*_b_ ≈ 0.04 s μm^-1^, and DDM measures 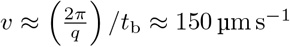, which is realistic for bull spermatozoa [34]. The third process collapses when *f*(*q, τ*) is plotted against *q*^2^*τ*, Fig. 4(b) (inset), indicating that it is diffusive, and therefore could come from the motion of non-motile spermatozoa and debris. The ISF of a diffusive process is a single exponential, with a characteristic time scale given by the decay of its amplitude to *e*^-1^ of its original value, which occurs at *q*^2^*t*_3_ ≈ 0.3 μm^-2^ s, giving an associated diffusivity *D* = 1/(*q*^2^*t*_3_) ≈ 3 μm^2^ s^-1^.

**Fig 4.**
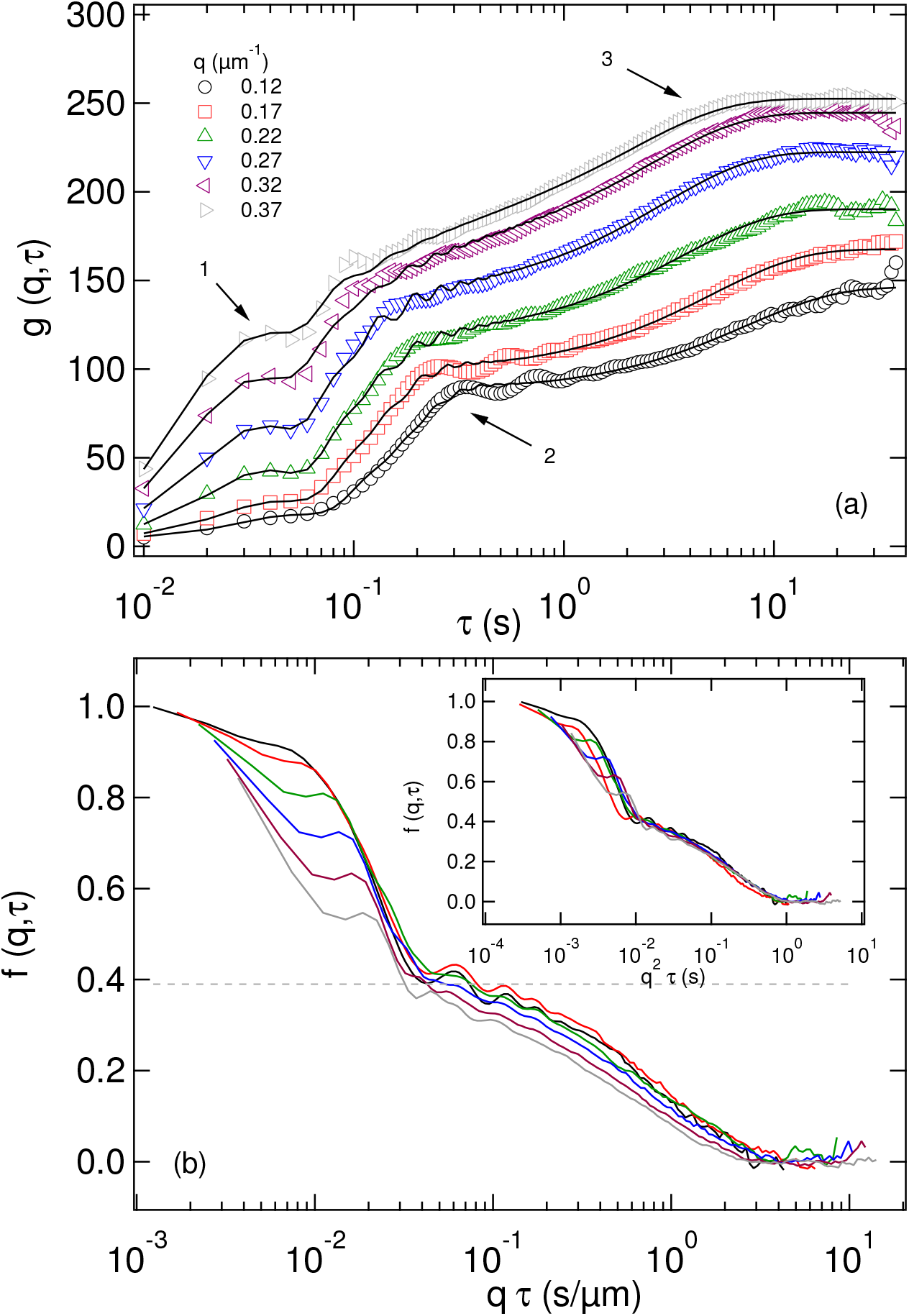
**DDM results** from a movie (2.5×, 100fps, 480p, 40s) of a diluted sample. (a) Measured *g*(*q, τ*) at 6 values of *q*, specified in the legend. Black lines are fits to the model given by Eq. 2. The arrows indicate three processes associated with head oscillation (1), swimming (2), and diffusion of non-motile sperm cells (3). (b) Reconstructed *f*(*q, τ*) plotted against *qτ* and *q*^2^*τ* (inset).

To help interpret this diffusivity and as a first step towards fitting measured *g*(*q, τ*)s, we left a fourfold-diluted thawed sample at room temperature until all motility ceased. Figure 5 shows the *g*(*q, τ*) at the highest measured *q* = 2.22μm^-1^ obtained from a movie of such a sample containing only non-motile spermatozoa and debris. Fitting a double exponential to this data (i.e., Eq. 8 in Eqs. 4 and 5 and assuming *α* = 0), we obtain two diffusivities: *D*_1_ = 0.018 ± 0.004 μm^2^ s^-1^ and *D*_2_ = 0.38 ± 0.06 μm^2^ s^-1^ (see bottom right inset Fig. 5).

**Fig 5.**
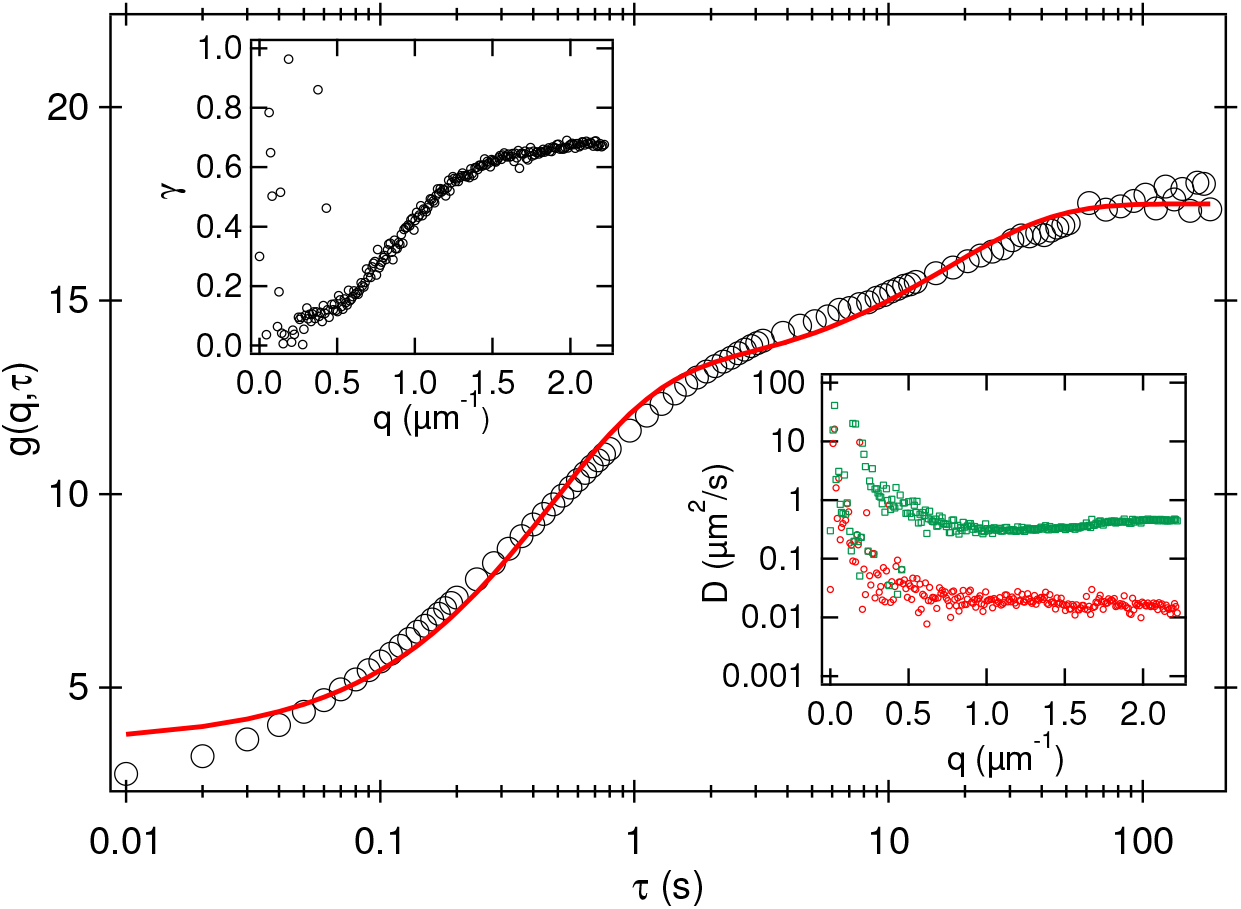
**Measured** *g*(*q, τ*) from a movie (10×, 100 fps, 500 p, 100s) of an in-active sample at *q* = 2.22*μ*m^-1^ with cell density of ≈ 20 × 10^6^cells/ml. Line is a fit using two exponential functions returning two separate diffusion coefficient *D*_1_ and *D*_2_. Right inset: fitted parameters *D*_1_ and *D*_2_ for a range of *q*. Left inset: Fitted measurement of the proportional contribution, *γ*, of the debris to the total signal.

The free diffusivity of a sphere with radius *R* ~ 5 μm, comparable to the head of a bull spermatozoa, is ~ 0.04 μm^2^ s^-1^, so that we identify *D*_1_ with *D*_nm_ for non-motile spermatozoa. We then reanalysed a cropped movie containing no (non-motile) sperm cells, and found a single process (data not shown). Fitting the measured *g*(*q, τ*) to a single exponential, yielding a diffusivity of 0.32 ± 0.06 μm^2^ s^-1^, so that *D*_2_ is *D*_d_ for debris. The fitting in fig. 5 also yielded the amplitudes of the contributions from (non-motile) spermatozoa and debris, *A*_s_ and *A*_d_, respectively. The ratio 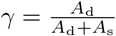, Fig. 5 (upper left inset), shows that non-motile spermatozoa dominate at low *q* (larger length scales), *γ* ≲ 0.1, while debris contribute to the signal at higher *q* (smaller length scales), *γ* ≳ 0.6.

Importantly, *γ* is independent of whether spermatozoa are motile or non-motile (see discussion around Eqs. 2 and 3). Returning to the data in Fig. 4, we therefore conclude that at these low *q* values, signal from spermatozoa dominate over signal from debris by a factor of 10 or more. The third, diffusive, process must therefore be associated with non-motile sperm cells. Visually, it is clear that this process contributes ≈ 40% of the amplitude, so that we conclude that the fraction of motile spermatozoa in this sample is *α* ≈ 0.6, comparable to the *α* = 0.66 ± 0.05 obtained by manual counting. The diffusivity of ~ 3 μm^2^ s^-1^ we estimated for non-motile spermatozoa in this sample is, however, ~ 100 × higher than the non-motile spermatozoa diffusivity measured from the sample shown in Fig. 5. This is due to the enhancement of passive diffusion by swimmers. The analogous effect in bacterial baths is well known [35, 36]. More importantly, enhancement of tracer diffusivity by an order of magnitude has been observed in suspensions of motile algae that swim at comparable speeds to our sperm cells but whose flagella are shorter [37]. This enhancement brings the diffusion of non-swimmers into a convenient time window (compare the time axes of Figs. 4 and 5) for measuring *α*.

### Motility parameters from data fitting

Since debris contribute ≲ 10% to the amplitude of the ISF at low *q* values, we take *A*_d_ = 0, and fitted the data shown in Fig. 4 to Eqs. 4, 5, 7 and 8 at each *q* to give *υ*(*q*), *σ*(*q*), *α*(*q*), *A*_0_(*q*), *f*_0_(*q*) and *D*_nm_(*q*), Fig. 6 (red data points). All fitted parameters except *D*_nm_ are approximately independent of *q* in the mid-range of *q* values shown. The precise window over which a parameter can be expected to be *q*-independent, and therefore can be meaningfully averaged over, depends on the physics of the associated process.

**Fig 6.**
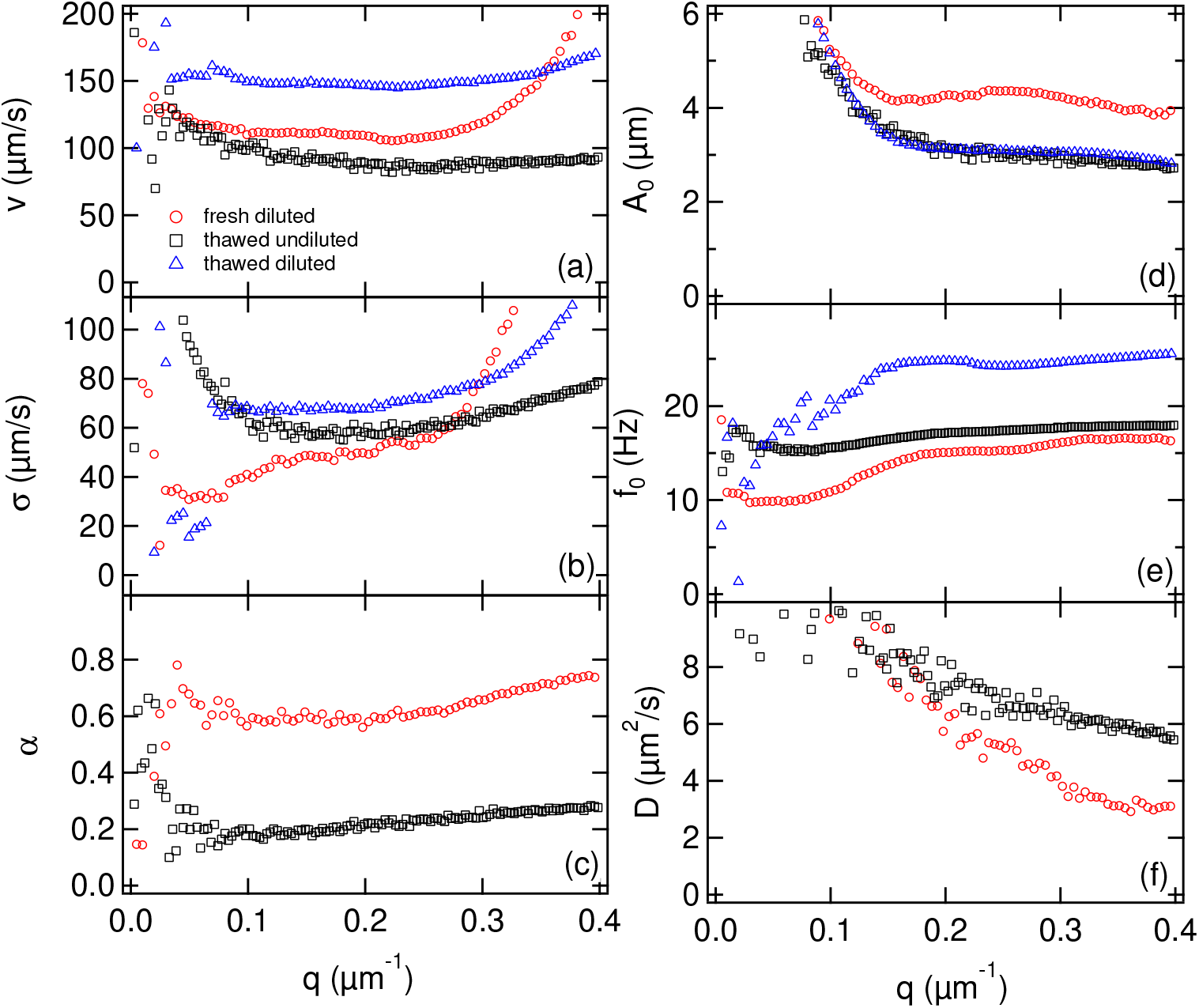
**Fitted parameters** over the range *q* = 0-0.4 μm^-1^. (a) Mean *υ* and (b) width *σ* of the fitted Schultz swimming speed distribution, (c) proportion of motile cells *α*, (d) head amplitude *A*_0_, (e) head frequency *f*_0_ and (f) diffusion coefficient of the non-motile spermatozoa *D*_nm_. (o) Fresh ejaculate diluted to ~ 20 × 10^6^ cells/ml and recorded using (2.5×, 100 fps, 480 p, 40 s). These parameters correspond to the data in Fig. 4. (□) thawed semen undiluted (~ 80 × 10^6^ cells/ml) and recorded using (2×, 300 fps, 300p, 100s). (Δ) thawed semen diluted 4× to ~ 20 × 10^6^ cells/ml recorded using (2×, 1000fps, 300p, 8s).

Characterising the oscillatory head motion is principally limited by low signal at low *q*, because this contribution to the ISF scales as *qA*_0_ (see Eq. 7). Thus, we find that *A*_0_(*q*) and *f*_0_(*q*) only become relatively constant at and beyond *q* ≈ 0.16 μm^-1^, where the contribution from head oscillation rises above ~ 10%. Averaging over *q*_min_ = 0.16 – 0.39*μ*m^-1^ (*l* ~ 16 – 39*μ*m) returns *A*_0_ = 4.2 ± 0.2 μm and *f*_0_ = 16 ± 1 Hz.

To characterise swimming, there must be a finite time interval 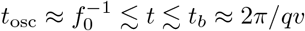, and so becomes problematic above *q*_max_ ≈ 2*πf*_0_/*υ* ≈ 0.5 μm^-1^ defined by the crossover in the characteristic time of the head oscillation and swimming processes, i.e. when *t*_osc_ = *t_b_*. In practice, fitted swimming parameters become unreliable before *q*_max_ is reached, signalled in our case by *υ* and *σ* becoming *q*-dependent above ≈ 0.3 μm^-1^. Averaging over *q* = 0.05-0.30μm^-1^ (*l* ~ 21-126μm) returns *υ* = 111 ± 3 μms^-1^, *σ* = 48 ± 9μms^-1^ and *α* = 0.61 ± 0.03.

In our *q* window, *D*_nm_ never becomes constant for this sample, although the data suggests that *D*_nm_ may become constant at *q* ≳ 0.3 μm^-1^. For the lowest *q*, this is perhaps partly because *f*(*q, τ*) has barely completed its decay in our time window (cf. Fig. 4(a)). More importantly, we know that the motion of non-motile organisms is enhanced by the presence of swimmers: the fitted values of *D*_nm_ are ~ 10^2^ × the thermal diffusivity measured from samples without motile cells (Fig. 5). There is no *a priori* reason to believe that it should be possible to model this motion as diffusive. In fact, doing so produces good fits, Fig. 4(a), but with a *q*-dependent diffusivity that is larger at larger length scales. It is inconsequential for our purposes that the physical origins of this effect are currently unknown, because empirically a a*q*-dependent *D*_nm_ produces good data fitting and gives correct motility parameters for the sperm cells (see next section).

What does matter is that the diffusivity of non-motile cells is enhanced from ~ 0.02 μm^2^ s^-1^, Fig. 5, to ≳ 4 μm^2^ s^-1^ by motile cells. To reliably measure *α* down to a *q*_min_ ≈ 0.1 μm^-1^ from the relative amplitudes of the active (swimming/head beating) and passive (non-motile diffusing) processes requires a time window of at least (*D*_nm_*q*^2^_min_)^-1^. An unenhanced *D*_nm_ would necessitate prohibitively long data acquisition times (≳ 1 h).

Note that we have fitted our data by assuming that the swimming speed distribution is single-peaked. The possibility of twin-peaked distributions is discussed in S2 Fig, where we also offer some comments on how to treat cases where a high proportion of spermatozoa swim in tight circles (see S3 Fig).

### In-lab validation of DDM

Figure 6 also shows DDM motility parameters extracted from fitting *g*(*q, τ*)’s obtained from movies recorded in-lab of thawed straws undiluted (~ 80 × 10^6^ cell/ml, black points) and diluted (~ 20 × 10^6^ cell/ml, blue points). The fitted parameters in both cases are constant over a greater *q* range than those obtained from the on-farm sample. This is due to either an increased frequency *f*_0_ (diluted) or a decreased speed *υ* (undiluted), thus extending *q*_max_. In the latter case, we cannot see the decay of the correlation function in the timescale of the movie (8s), and therefore have no measure of *α* or *D*_nm_ in this case. This was consistently true for dilute, thawed samples as the diffusivity of their non-motile cells was less enhanced than in undiluted samples with a higher concentration or fresh samples with a higher motile fraction.

To validate DDM for measuring bull spermatozoa motility parameters, we compared DDM to particle tracking in the laboratory. The Schultz distributions obtained from fitting the *g*(*q, τ*)s of a sample at two different times are compared to the histogram of swimming speeds (VAP and VSL) calculated from tracking in Fig. 7. Note that in DDM analysis, non-motile cells (which, in practice, includes all tracked trajectories with 0 ≤ *υ* ≤ 20 μms^-1^) are not included in the Schultz *P*(*υ*), but are separately accounted for in terms of the non-motile fraction, (1 – *α*). Taking this into account, we find that the DDM swimming speed distribution is consistent with the tracked distribution of either VAP or VSL. We do not expect *P*(VAP) and *P*(VSL) to differ greatly over the time window of our movie (0.5 s) because the swimming tracks have low curvature on this time scale. Inspection of ~ 50 tracks of swimming spermatozoa returned values for *A*_0_ and *f*_0_ that agree with DDM measurements, Fig. 7, and are consistent with values obtained from CASA in previous studies [6,38].

**Fig 7.**
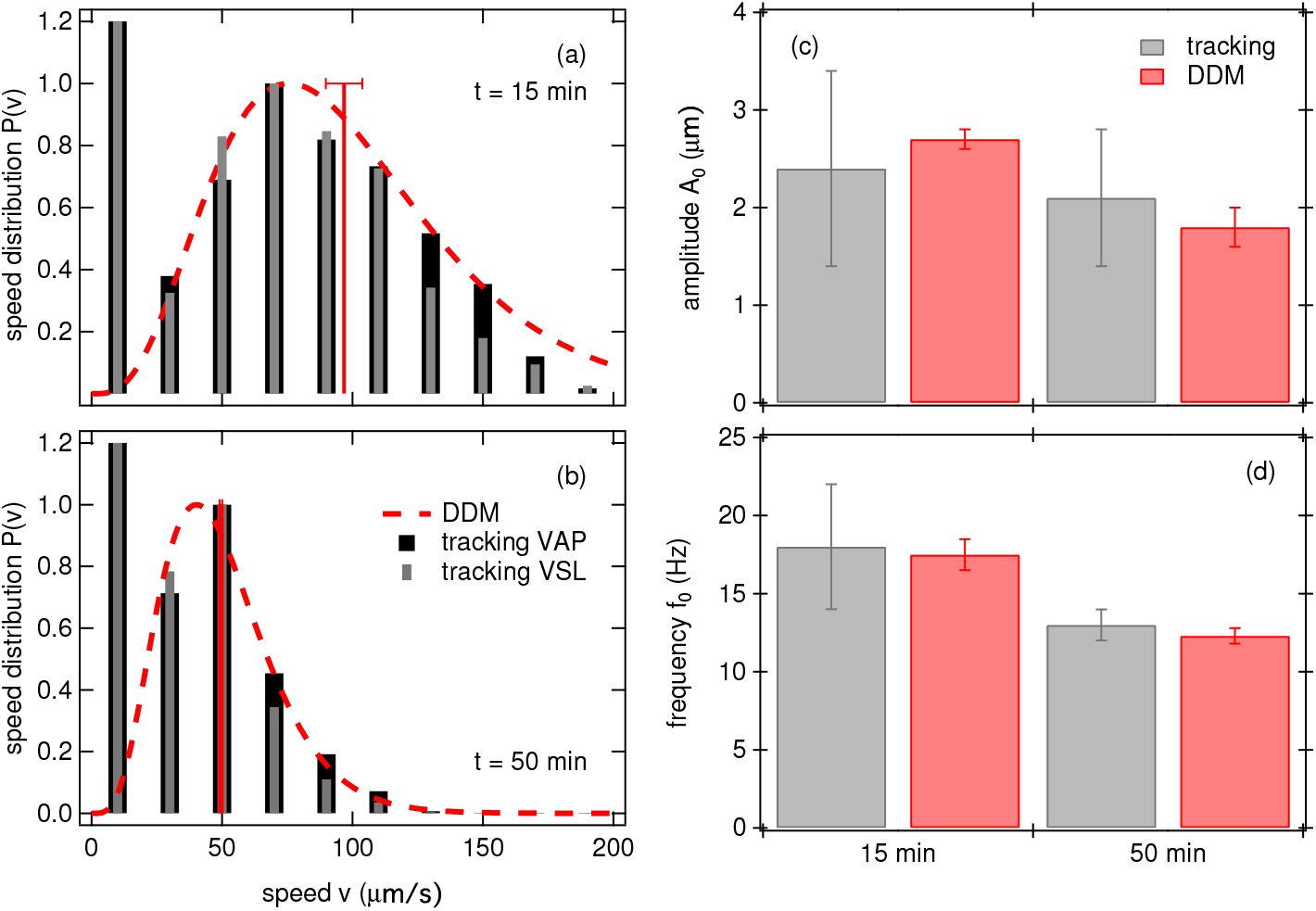
**A comparison of tracking and DDM methods** for a sample maintained at 37°C for 15 min and 50 min after thawing. (a,b) swimming speed distributions. Histogram of VAP (black) and VSL (grey) were calculated from 10 movies (10×, 100fps, 500p, 0.5s) with ≈ 50 swimming tracks per movie. A movie (300fps, 500p, 10s) at 2× magnification was recorded immediately afterwards and analysed with DDM to give *υ* (red vertical bar) and *σ* from which *P*(*υ*) was reconstructed (red dashed line). Histograms are normalised in order that the peak is at 1. (c,d) Head oscillation amplitude *A*_0_ and frequency *f*_0_ measured with DDM and tracking for consecutive movies of the same sample. Note that the quoted error for tracking is the standard deviation for measurements from 50 tracks, while that for DDM is a standard deviation of the mean for averaging over the values measured at a set of *q* values.

To validate the measurement of *α*, we studied samples containing sufficient motile cell density to enhance the diffusivity of non-motile cells so that the latter contributes to the ISF in our time and *q* window. Table 1 compares the motile fraction measured from direct counting and from DDM, again showing agreement.

**Table 1.** Motile fraction: DDM vs tracking.

| | $\alpha_{\text{DDM}}$ (%) | $\alpha_{\text{Counting}}$ (%) | $v$ ($\mu\text{m s}^{-1}$) | $A_0$ ($\mu\text{m}$) | $f_0$ (Hz) |
| --- | --- | --- | --- | --- | --- |
| sample 1 | $61 \pm 3$ | $66 \pm 5$ | $111 \pm 3$ | $4.2 \pm 0.2$ | $16.0 \pm 1$ |
| sample 2 | $25 \pm 3$ | $22 \pm 5$ | $107 \pm 1$ | $4.0 \pm 0.2$ | $13.6 \pm 0.2$ |
| sample 3 | $42 \pm 5$ | $43 \pm 5$ | $82 \pm 2$ | $3.2 \pm 0.2$ | $14.0 \pm 0.5$ |
| sample 4 | $15 \pm 5$ | $18 \pm 5$ | $76 \pm 5$ | $2.6 \pm 0.1$ | $17.0 \pm 1$ |

### Relationship between *υ*, *A*_0_ and *f*_0_

To demonstrate the potential of DDM for fundamental research in spermatozoa motility, we use the method to verify a basic theoretical result. Since Taylor’s pioneering work [39], the motility of flagellated microorganisms at low Reynolds number has been studied in detail. Thus, e.g., Keller and Rubinow found [40] that a spherical body joined to an elastic filament (the flagellum) of length *L* performing planar or helical wave motion [40] is propelled at 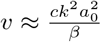, where *c*, *λ* = 2*π*/*k* and *a*_0_ are the speed, wavelength and amplitude of the undulations propagating along the flagellum, or, in terms of the frequency 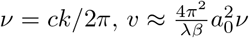. *β* depends on the shape and motion of the sperm cells and is a function of the geometry of both head an flagellum. Importantly, if *a*_0_/*L* → 0, the free end of the flagellum exerts no torque on the head, and frequency and amplitude of body and flagellum become the same: *f*_0_ → *ν*, *A*_0_ → *a*_0_, so that

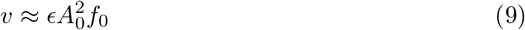

with *ϵ* = 4*π*^2^/*λβ*. Analytical expressions for *β* are given in [40] for planar, 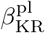, and helical motion, 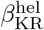 (see S4 text). Although *β* displays a weak dependency with (*kA*_0_)^2^, we expect *β* to be approximately constant over the typical range of values for *A*_0_ and *λ*, and thus 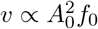. Note that the 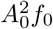 scaling has been predicted by others, but with different prefactors [41–44].

We explored the validity of this relationship by monitoring (*υ, A*_0_, *f*_0_) over 120 min in three independent, thawed undiluted samples (see caption of Fig. 8 for details). These parameters changed with time, especially as the cells gradually depleted the suspending medium of oxygen [45, 46]. The three samples monitored over time therefore gave a range of these parameters: *υ* = 50-200μms^-1^, *A*_0_ = 2-4 μm and *f*_0_ = 5-20Hz. All data at speeds above ~ 60*μ*m/s collapse onto a universal curve when *υ* is plotted against 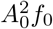, Fig. 8 thus confirming the approximate scaling of Eq. 9. A linear fit through the origin gives the prefactor, which varies marginally between samples, *ϵ*_exp_ = 0.52 – 0.62 μm^-1^, suggesting slight variability in spermatozoa morphology. Interestingly, we found our experimental prefactor to be closer to 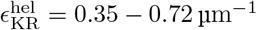 than 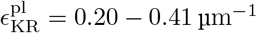 predicted for helical and planar motion respectively, assuming typical bull sperm head radius (4 μm), flagellar radius (0.4μm), length (40 – 60 μm), wavelength (λ = 30 – 60 μm), and measured amplitude *A*_0_ = 2 – 4 μm. This suggests that sperm flagella follow predominantly a helical motion rather than planar motion in the present experiments. Indeed, a previous study has identified that flagellum follows a planar wave mode when both body and flagellum are confined to within 1 μm of a wall [47]. In our present experiments, we image cells swimming through the whole height 20 μm-chamber and thus expect cells not strictly confined to within 1 μm of the wall.

**Fig 8.**
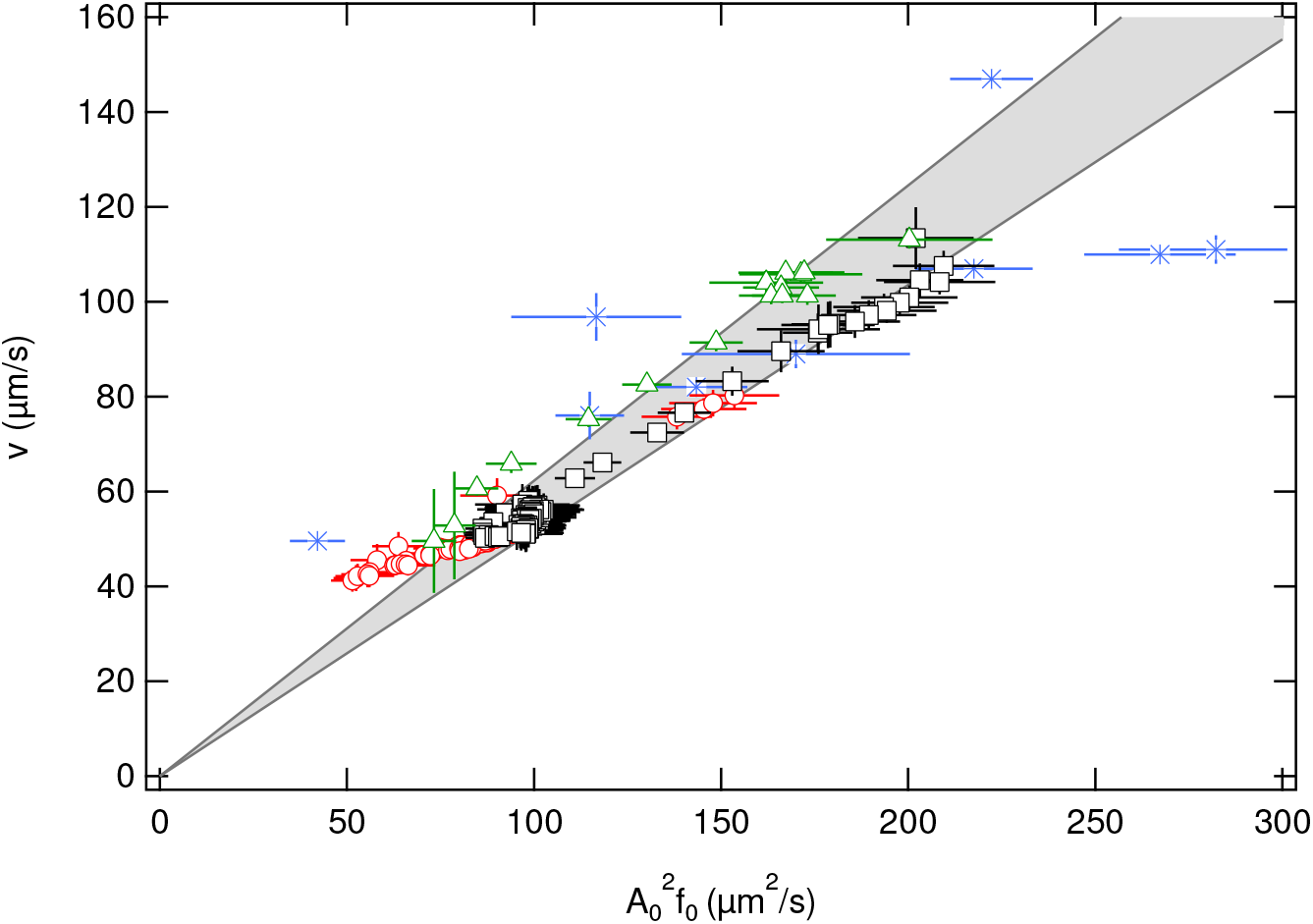
**Scaling.** *υ* plotted against 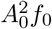 for DDM measurements of thawed straws: BB (□), CH (Δ) and HO semen (∘) were pipetted undiluted into pre-warmed sample chambers immediately (≈ 80 × 10^6^cells/ml). Movies (2×, 300fps, 300p, 40s) were recorded every 2 min subsequently. (Only the first 30 min of CH are included, before the data becomes noisy.) Data from Fig. 6, Fig. 7 and Table 1 plotted in (*). Grey area defines the range of linear fit through the origin for all three dataset and 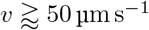.

## Conclusion

We have demonstrated that DDM is a high-throughput technique for characterising bull semen motility that can be applied to both fresh samples on farm and thawed samples in the laboratory. The technique was verified with particle tracking, the current method of choice in veterinary practice.

Currently bull pre-breeding examination includes subjective visual microscopy. CASA provides objective measurements in-lab but is rarely used on-farm. DDM could offer an objective bull-side evaluation in pre-breeding examinations and evaluations of thawed AI semen on-farm. Additionally, measurements could be performed on samples whose analysis is problematic with CASA, e.g. spermatozoa in milk extender. The portability and speed of our technique should enable large-scale studies to correlate motility parameters with field fertility outcomes, thus providing evidence-based guidelines for the interpretation of data collected by other methods such as CASA and flow cytometry [48].

The high-throughput nature of our technique enabled us to collect a large enough data set in the laboratory to verify a long-standing theoretical prediction relating different kinematic parameters of spermatozoa motion. This demonstrates the utility of DDM as a high-statistic method of assessing motility in varying environmental conditions – in our case probably progressive oxygen depletion. The technique can easily be automated, and applied to the study of different sample geometries such as confinement [49,50]. Our method is not in principle restricted to bull semen, and can be used to study the correlation between motility and fertility as well as the physiology of spermatozoon swimming. Additionally, it should also impact fundamental research in biophysics and active matter physics, where various aspects of spermatozoon swimming attract attention.

## Acknowledgement

We thank Drs Martin Tomlinson, Isobel Truyers, Paul Wood and Fraser Murdoch from the Royal (Dick) School of Veterinary Studies at the Roslin Institute (University of Edinburgh) for allowing us to attend their visits to farms in order to measure fresh samples, and Dr Alexander N. Morozov for helpful discussions. The work was funded by the UK EPSRC (EP/K503794/1 and EP/J007404/1) and the European Research Council (AdG 340877-PHYSAPS and PoC 72774-PorCSperM).

## Supporting information

**S1 Fig. Counting motile spermatozoa.**

**S2 Fig. Special case: two speed distributions.**

**S3 Fig. Special case: circular motion.**

**S4 text. Analytical expression for *β* in Eq. 9 of main text.**

## References

1. Ruestow EG. Images and ideas: Leeuwenhoek’s perception of the spermatozoa. J Hist Biol. 1983;16(2):185–224. doi:10.1007/BF00124698.

2. Humphries S, Evans JP, Simmons LW. Sperm competition: linking form to function. BMC Evol Biol. 2008;8.

3. Baccetti B. Evolution of the spermatozoon. Bolletino di zoologia. 1984;51:25–33.

4. Poiani A. Complexity of seminal fluid: a review. Behav Ecol Sociobiol. 2006;60:289–310.

5. Davis RO, Katz DF. Operational standards for CASA instruments. J Androl. 1993;14(5):385–94.

6. Amann RP, Waberski D. Computer-assisted sperm analysis (CASA): capabilities and potential developments. Theriogenology. 2014;81(1):5–17.e1–3. doi:10.1016/j.theriogenology.2013.09.004.

7. Elgeti J, Kaupp UB, Gompper G. Hydrodynamics of sperm cells near surfaces. Biophys J. 2010;99(4):1018–26. doi:10.1016/j.bpj.2010.05.015.

8. Martinez-Pastor F, Tizado EJ, Garde JJ, Anel L, de Paz P. Statistical Series: Opportunities and challenges of sperm motility subpopulation analysis; 2011.

9. Boyers SP, Davis RO, Katz DF. Automated semen analysis. Curr Probl Obstet Gynecol Fertil. 1989;12:167–200.

10. Cerbino R, Trappe V. Differential Dynamic Microscopy: Probing Wave Vector Dependent Dynamics with a Microscope. Phys Rev Lett. 2008;100(18):188102. doi:10.1103/PhysRevLett.100.188102.

11. Wilson LG, Martinez VA, Schwarz-Linek J, Tailleur J, Bryant G, Pusey PN, et al. Differential Dynamic Microscopy of Bacterial Motility. Phys Rev Lett. 2011;106(1):018101. doi:10.1103/PhysRevLett.106.018101.

12. Martinez VA, Besseling R, Croze OA, Tailleur J, Reufer M, Schwarz-Linek J, et al. Differential dynamic microscopy: a high-throughput method for characterizing the motility of microorganisms. Biophys J. 2012;103(8):1637–47. doi:10.1016/j.bpj.2012.08.045.

13. Kastelic J, Thundathil J. Breeding Soundness Evaluation and Semen Analysis for Predicting Bull Fertility. Reprod Dom Anim. 2008;43:368–373.

14. Condorelli RA, La Vignera S, Bellanca S, Vicari E, Calogero AE. Myoinositol: does it improve sperm mitochondrial function and sperm motility? Urology. 2012;79(6):1290–5. doi:10.1016/j.urology.2012.03.005.

15. Publicover SJ, Barratt CLR. Sperm motility: things are moving in the lab! Mol Hum Reprod. 2011;17(8):453–6. doi:10.1093/molehr/gar048.

16. Suarez SS. Control of hyperactivation in sperm. 2008;14(6):647–657. doi:10.1093/humupd/dmn029.

17. Friedrich BM, Jülicher F. Chemotaxis of sperm cells. Proc Natl Acad Sci USA. 2007;104(33):13256–13261. doi:10.1073/pnas.0703530104.

18. Kaupp UB, Kashikar ND, Weyand I. Mechanisms of sperm chemotaxis. Annu Rev Physiol. 2008;70:93–117. doi:10.1146/annurev.physiol.70.113006.100654.

19. Jikeli JF, Alvarez L, Friedrich BM, Wilson LG, Pascal R, Colin R, et al. Sperm navigation along helical paths in 3D chemoattractant landscapes. Nature Comm. 2015;6:7985. doi:10.1038/ncomms8985.

20. Riedel IH, Kruse K, Howard J. A self-organized vortex array of hydrodynamically entrained sperm cells. Science. 2005;309(5732):300–3. doi:10.1126/science.1110329.

21. Gaffney EA, Gadêlha H, Smith DJ, Blake JR, Kirkman-Brown JC. Mammalian Sperm Motility: Observation and Theory. Annu Rev Fluid Mech. 2011;43(1):501–528. doi:10.1146/annurev-fluid-121108-145442.

22. Immler S, Moore HDM, Breed WG, Birkhead TR. By hook or by crook? Morphometry, competition and cooperation in rodent sperm. PLoS one. 2007;2(1):e170. doi:10.1371/journal.pone.0000170.

23. Kantsler V, Dunkel J, Blayney M, Goldstein RE. Rheotaxis facilitates upstream navigation of mammalian sperm cells. eLife. 2014;3(3):e02403. doi:10.7554/eLife.02403.001.

24. Denissenko P, Kantsler V, Smith DJ, Kirkman-Brown J. Human spermatozoa migration in microchannels reveals boundary-following navigation. Proc Natl Acad Sci USA. 2012;109(21):8007–8010. doi:10.1073/pnas.1202934109.

25. Guidobaldi A, Jeyaram Y, Berdakin I, Moshchalkov VV, Condat CA, Marconi VI, et al. Geometrical guidance and trapping transition of human sperm cells. Physical Review E. 2014;89(032720):1–6. doi:10.1103/PhysRevE.89.032720.

26. Guidobaldi HA, Jeyaram Y, Condat CA, Oviedo M, Berdakin I, Moshchalkov VV, et al. Disrupting the wall accumulation of human sperm cells by artificial corrugation. biomicrofluidics. 2015;9(024122):1–9.

27. Schwarz-Linek J, Arlt J, Jepson A, Dawson A, Vissers T, Miroli D, et al. Escherichia coli as a model active colloid: A practical introduction. Colloids Surf B. 2016;137:2–16. doi:10.1016/j.colsurfb.2015.07.048.

28. Hallett FR, Craig T, Marsh J. Swimming speed distributions of bull spermatozoa as determined by quasi-elastic light scattering. Biophys J. 1978;21(3):203–16. doi:10.1016/S0006-3495(78)85520-9.

29. Craig T, Hallett FR, Nickel B. Quasi-elastic light-scattering spectra of swimming spermatozoa. Rotational and translational effects. Biophys J. 1979;28(3):457–72. doi:10.1016/S0006-3495(79)85193-0.

30. Craig T, Hallett FR, Nickel B. Motility analysis of circularly swimming bull spermatozoa by quasi-elastic light scattering and cinematography. Biophys J. 1982;38(1):63–70. doi:10.1016/S0006-3495(82)84531-1.

31. Shimizu H, Matsumoto G. Light scattering study on motile spermatozoa. IEEE Trans Biomed Eng. 1977;24(2):153–7. doi:10.1109/TBME.1977.326120.

32. Schneider CA, Rasband WS, Eliceiri KW. NIH Image to ImageJ: 25 years of image analysis. Nature Methods. 2012;9(7):671–675.

33. Crocker JC, Grier DG. Methods of Digital Video Microscopy for Colloidal Studies. J Colloid Interface Sci. 1996;179(1):298–310. doi:10.1006/jcis.1996.0217.

34. Kathiravan P, Kalatharan J, Karthikeya G, Rengarajan K, Kadirvel G. Objective Sperm Motion Analysis to Assess Dairy Bull Fertility Using Computer-Aided System - A Review. Reprod Domest Anim. 2011;46(1):165–172. doi:10.1111/j.1439-0531.2010.01603.x.

35. Miño G, Mallouk TE, Darnige T, Hoyos M, Dauchet J, Dunstan J, et al. Enhanced Diffusion due to Active Swimmers at a Solid Surface. Phys Rev Lett. 2011;106:048102+. doi:10.1103/physrevlett.106.048102.

36. Jepson A, Martinez VA, Schwarz-Linek J, Morozov A, Poon WCK. Enhanced diffusion of nonswimmers in a three-dimensional bath of motile bacteria. Phys Rev E. 2013;88(4):041002. doi:10.1103/PhysRevE.88.041002.

37. Leptos KC, Guasto JS, Gollub JP, Pesci AI, Goldstein RE. Dynamics of Enhanced Tracer Diffusion in Suspensions of Swimming Eukaryotic Microorganisms. Phys Rev Lett. 2009;103:198103.

38. Muiño R, Tamargo C, Hidalgo CO, Peña AI. Identification of sperm subpopulations with defined motility characteristics in ejaculates from Holstein bulls: Effects of cryopreservation and between-bull variation. Anim Reprod Sci. 2008;109(1-4):27–39. doi:10.1016/j.anireprosci.2007.10.007.

39. Taylor G. The Action of Waving Cylindrical Tails in Propelling Microscopic Organisms. Proc Royal Soc Lond A. 1952;211(1105).

40. Keller JB, Rubinow SI. Swimming of flagellated microorganisms. Biophys J. 1976;16:151–170.

41. Gray J, Hancock GJ. The propulsion of sea-urchin spermatozoa. J Exp Biol. 1955;32:802.

42. Lighthill MJ. Mathematical Biofluiddynamics. SIAM, Philadelphia; 1975.

43. Lauga E. Floppy swimming: Viscous locomotion of actuated elastica. Phys Rev E. 2007;75(4):041916. doi:10.1103/PhysRevE.75.041916.

44. Friedrich BM, Riedel-Kruse IH, Howard J, Julicher F. High-precision tracking of sperm swimming fine structure provides strong test of resistive force theory. J Exp Biol. 2010;213(8):1226–1234. doi:10.1242/jeb.039800.

45. Wilson MC, Harvey JD, Shannon P. Aerobic and anaerobic swimming speeds of spermatozoa investigated by twin beam laser velocimetry. Biophys J. 1987;51(3):509–512. doi:10.1016/S0006-3495(87)83373-8.

46. Vishwanath R, Shannon P. Do sperm cells age? A review of the physiological changes in sperm during storage at ambient temperature. In: Reproduction, Fertility and Development. vol. 9; 1997. p. 321–331.

47. Nosrati R, Driouchi A, Yip CM, Sinton D. Two-dimensional slither swimming of sperm within a micrometre of a surface. Nature Comm. 2015;6:8703. doi:10.1038/ncomms9703.

48. Sellem E, Broekhuijse MLWJ, Chevrier L, Camugli S, Schmitt E, Schibler L, et al. Use of combinations of in vitro quality assessments to predict fertility of bovine semen. Theriogenology. 2015;84(9):1447–1454.e5. doi:10.1016/j.theriogenology.2015.07.035.

49. Contri A, Valorz C, Faustini M, Wegher L, Carluccio A. Effect of semen preparation on casa motility results in cryopreserved bull spermatozoa. Theriogenology. 2010;74(3):424–35. doi:10.1016/j.theriogenology.2010.02.025.

50. Gee C, Zimmer-Faust R. The effects of walls, paternity and ageing on sperm motility. J Exp Biol. 1997;200(24):3185–3192.

